# Shannon: An Information-Optimal de Novo RNA-Seq Assembler

**DOI:** 10.1101/039230

**Authors:** Sreeram Kannan, Joseph Hui, Kayvon Mazooji, Lior Pachter, David Tse

**Affiliations:** University of Washington, Seattle.; University of California, Berkeley; University of California, Los Angeles; Stanford University and University of California, Berkeley

## Abstract

De novo assembly of short RNA-Seq reads into transcripts is challenging due to sequence similarities in transcriptomes arising from gene duplications and alternative splicing of transcripts. We present Shannon, an RNA-Seq assembler with an optimality guarantee derived from principles of information theory: Shannon reconstructs nearly all information-theoretically reconstructable transcripts. Shannon is based on a theory we develop for de novo RNA-Seq assembly that reveals differing abundances among transcripts to be the key, rather than the barrier, to effective assembly. The assembly problem is formulated as a sparsest-flow problem on a transcript graph, and the heart of Shannon is a novel iterative flow-decomposition algorithm. This algorithm provably solves the information-theoretically reconstructable instances in linear-time even though the general sparsest-flow problem is NP-hard. Shannon also incorporates several additional new algorithmic advances: a new error-correction algorithm based on successive cancelation, a multi-bridging algorithm that carefully utilizes read information in the k-mer de Bruijn graph, and an approximate graph partitioning algorithm to split the transcriptome de Bruijn graph into smaller components. In tests on large RNA-Seq datasets, Shannon obtains significant increases in sensitivity along with improvements in specificity in comparison to state-of-the-art assemblers.

---

De novo RNA-Seq assembly, in the absence of a reference genome, is an important first step for transcriptomic analysis of non-model organisms as well as in cancer transcriptomics (where the genome may be significantly disrupted). There have been numerous efforts to design de novo transcriptome assemblers, including SOAPdenovo-Trans,^1^ Oases,^2^ Trinity,^3^ and TransAbyss^4^; however de novo transcriptome assembly from short reads still remains a challenging problem due to the numerous similar isoforms of genes, the ubiquity of paralogs, varying abundances of transcripts, and the large numbers of reads required for assembly. For these reasons, de novo transcriptome assembly is considered hard and studies of assemblers reveal discordance in the output of methods (for example^5^).

In this letter, we describe Shannon, a new de novo RNA-Seq assembler designed using principles of information theory. In order to demonstrate the performance of Shannon and to understand the relative merits of Shannon with respect to state-of-the-art assemblers, we compared the performance of Shannon with four other de novo short-read assemblers: SOAPdenovo-Trans, TransABySS, Oases and Trinity, on two recently sequenced RNA-Seq datasets^a^. The datasets were specifically chosen because they include both “long” PacBio reads that can be used for validation, as well as deep (> 100 million) “short” Illumina reads which are input into the assemblers.

The first dataset comprises of 135M Illumina single-end reads, each 50 bp long, generated from an RNA sample from human embryonic stem cells.^6^ In Figure 1(a), we show the number of transcripts reconstructed accurately from an enhanced annotation. In Figure 1 (b) and (c), sensitivity of the various assemblers segregated by abundance and splicing complexity is shown, and in Figure 1(d), we show the specificity. We observe a uniform performance gain for Shannon over other assemblers. The second dataset^7^ comprises of 110 million Illumina 101-bp paired end reads from the Lymphoblastoid transcriptome in the GM12878 cell line. The performance is shown in Figures 1(d),(e), (f) and (g), and Shannon is seen to perform consistently better than other assemblers (Oases failed to run on this dataset).

**Figure 1:**
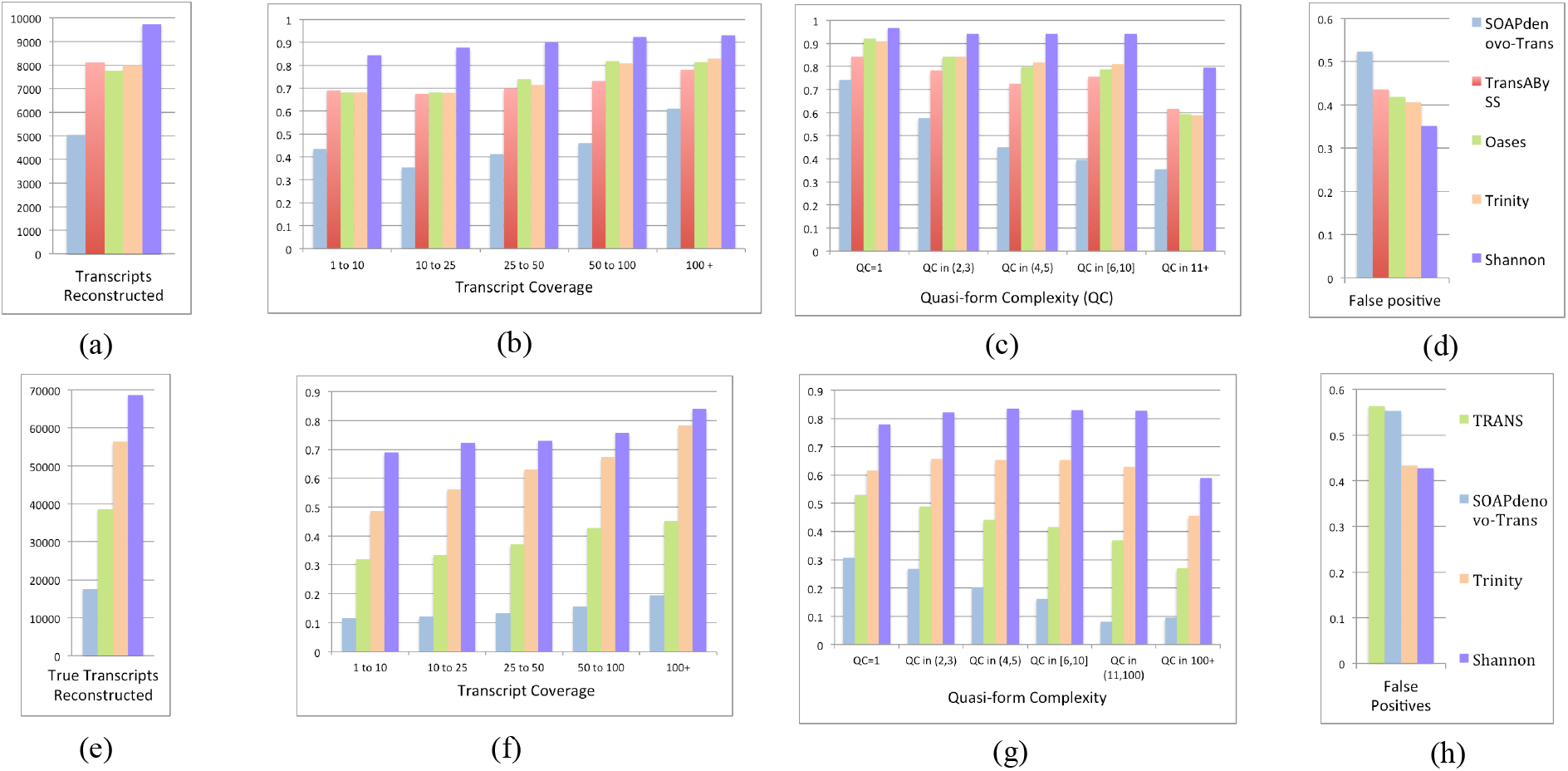
(a),(b),(c),(d): Results on a human embryonic stem cell transcriptome with 135 M single-end Illumina reads (50 bp). The “ground truth” used to evaluate sensitivity in (a) and (b) is obtained as follows. First, we take the 13,274 transcripts, which are assembled by the hybrid assembler IDP^6^ from the 135M short reads together with 7.8 M long PacBio reads. These transcripts are used to augment the RefSeq annotation to 54233 transcripts, out of which the number reconstructed accurately are shown in (a). A subset of 2885 of these transcripts which are fully covered by 25 length kmers from short reads; this is taken to be our “oracle set”^3^ and is used to evaluate recall rates of Shannon and other assemblers. We compare the fraction of transcripts recovered by different assemblers, segregated by coverage depths of the transcripts in (b), and segregated by splicing complexity of the transcripts in (c). We measure splicing complexity of a transcript by the size of the transcript family to which it belongs (two transcripts are in the same family if they share a kmer, of length 25). To measure false positive rates, we compare the reconstructed transcripts to the error-corrected long PacBio reads. We see in (d) that Shannon also performs best in terms of the false positive rate. (e),(f),(g),(h): Results on a dataset of 110 M paired-end short reads (101 bp) from Lymphoblastoid cells. Short reads are combined with long PacBio reads line (700K CCS reads) in order to augment the Gencode annotation to have 207266 transcripts.^7^ (e) Number of transcripts in enhanced Gencode reconstructed accurately. We select from the augmented Gencode annotation those transcripts which are fully covered by kmers (of length 25) in the Illumina reads to get the oracle set with 14186 transcripts used to measure sensitivity. (f) and (g) display the transcript reconstruction segregated by coverage depth and by splicing complexity respectively, and (h) displays the false positive rate measured against long reads. Shannon performs significantly better across both datasets across different categories.

We hypothesized that the significantly improved performance of Shannon in comparison to existing programs is due to the design of Shannon utilizing information theoretic principles rather than heuristics. Existing assemblers do not explicitly optimize performance metrics measuring reconstruction accuracy, and instead rely on ad-hoc heuristics that are guided by intuition. Specifically, we were unable to find any guarantees on reconstruction accuracy for any of the previously published assemblers we investigated. This lack of theoretical guarantees may be due in part to previous computer science work showing that various formulations of assembly are NP-hard.^8^, ^9^ We take a different view of the problem based on information theory, which turns out to neatly sidestep the NP-hardness barrier.

Information theory, originally invented in the context of communications,^10^ characterizes the minimum amount of data needed to solve an inference task. The theory was extended to DNA sequencing,^11^ where the minimum coverage depth and read length required to reconstruct a genome was calculated - see Figure 2 (a). An interesting conclusion from that paper is that, for any genome, there is a critical read length, called *l_crit_*, below which unique reconstruction is impossible at any coverage depth. Yet when read length is greater than *l_crit_*, Lander-Waterman coverage^12^ is essentially sufficient to reconstruct the genome. This critical read length is explicitly characterized in terms of certain bottleneck repeats in the genome. In this letter, we prove an analogous result for RNA-Seq assembly, providing a map of the fundamental information barriers faced by any assembly algorithm, while revealing an approach that can optimize assembly with respect to the inherent limits.

**Figure 2:**
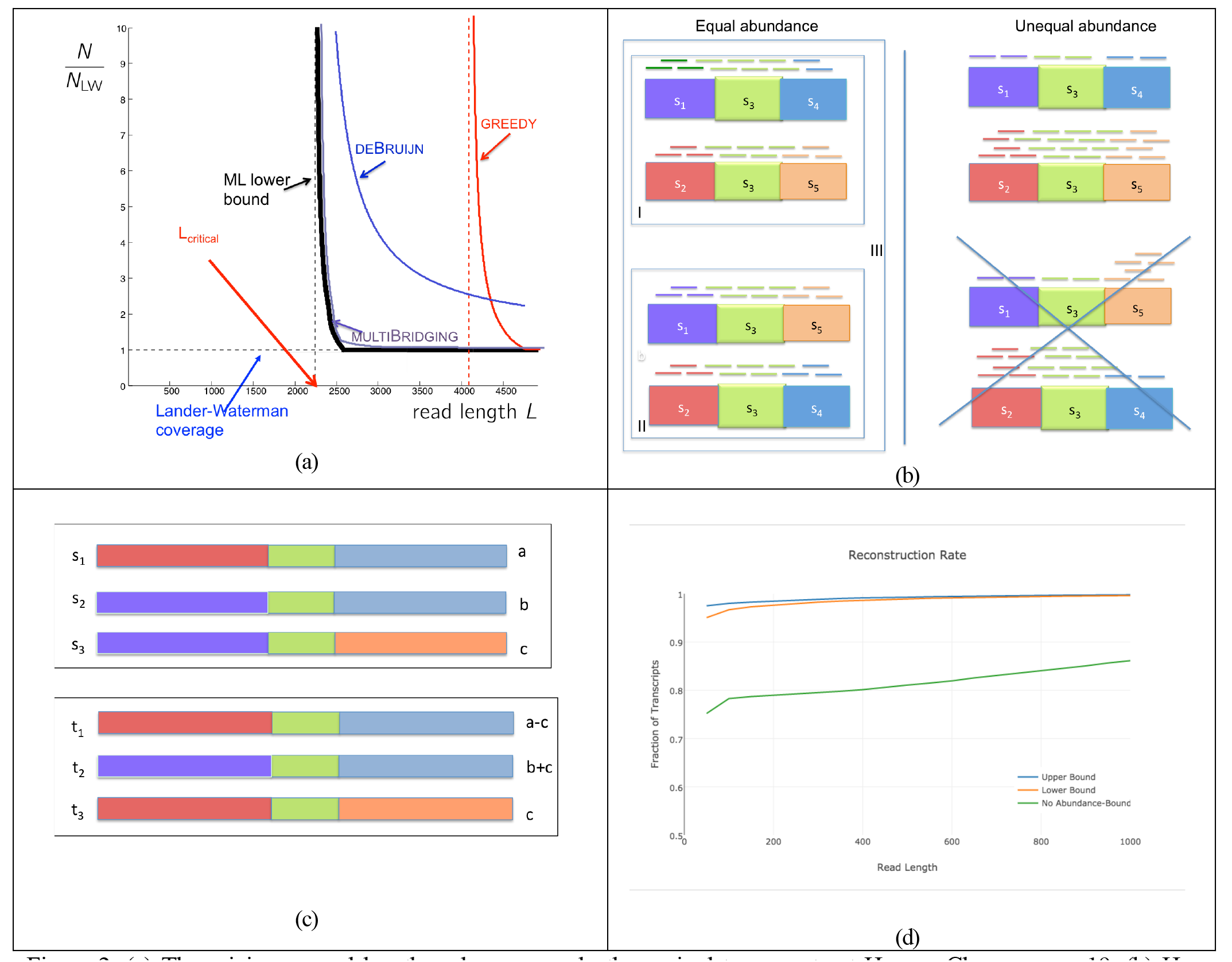
(a) The minimum read length and coverage depth required to reconstruct Human Chromosome 19. (b) How transcript abundances helps to disambiguate isoforms. The figure shows transcriptomes with isoforms sharing an exon s3 whose length is longer than the read length L. On the left hand side, we show the scenario where the abundances of all isoforms are the same. There are three possible transcriptomes consistent with the reads: (I) two isoforms, (II) two isoforms alternatively spliced, and (III) all four isoforms. On the right hand side, we consider the (more typical) scenario when the abundances of the isoforms are unequal. Here, there is only one unique transcriptome consistent with the read coverage statistics. The other solution (shown crossed out) will lead to non-uniform read coverage along each transcript. (c) A scenario in which isoforms cannot be disambiguated even with abundance information. In this example, the isoforms all share an exon (yellow region) of length greater than the read length L. There are two transcriptiomes consistent with the read coverage statistics. The first transcriptome has three isoforms with abundances a > b > c. The second transcriptome has three alternatively spliced isoforms with abundances a - c, b + c, c. Even though the abundances of the isoforms are different in the two transcriptomes, the frequencies of the reads from each of the colored regions are the same. Reads cannot bridge the yellow region while statistics are indistinguishable. (d) Theoretical bounds on the fraction of the HESC transcriptome that can be reconstructed for a given read length in the idealistic setting of infinite coverage depth and error-free reads. Upper curve: information theoretic upper bound on the largest fraction of the transcriptome that can be recovered. Middle: performance achieved by the proposed algorithm. Much below: performance bound for assembly algorithms that do not use abundance information to resolve repeats.

In the DNA assembly problem, long repeats within a genome are the barriers to reconstruction. In the RNA-Seq assembly problem, in addition to repeats inside transcripts, repeats across transcripts create a fundamental information barrier, which is typically the bottleneck. These inter-transcript repeats are common in higher Eukaryotes that are subject to alternative splicing; for example there may be multiple shared exons between different isoforms of genes. The left hand side of Figure 2(b) shows a case where multiple isoforms can give rise to the same reads and hence unique reconstruction is impossible. However, this example is problematic only in the special case when the transcripts have the exact same abundances. In generic cases, where transcripts have different abundances, it becomes possible to disambiguate between the two possible transcriptomes. Thus differing abundances is a key rather than a hurdle to unique transcriptome reconstruction - especially in resolving complex isoforms and paralogues. However, even with different abundances, certain complicated repeat structures in transcriptomes are impossible to reconstruct, see Figure 2(c) (more details in Supplementary Materials Section 3). By identifying these bottleneck repeat structures, we derived a transcriptome-dependent upper bound on the fraction of transcripts that can be reconstructed at any given read length; Figure 2(d) shows the limits for the HESC transcriptome, where no more than 95% of the transcripts can be reconstructed at read length 100 (upper curve). The middle curve shows that a new algorithm utilizing abundance information can result in performance close to the theoretical bound, achieving nearly 95% at the same read length. However, methods that do not appropriately leverage abundance information cannot reconstruct more than 80% of the transcripts accurately at read-length 100 (lower curve). See Online Methods for more details on the general theory for the derivation of these curves.

In the context of communications, information theory not only imposes a limit on how fast one can communicate but also leads to the design of efficient algorithms and codes to reach that limit. Analogously, the information theoretic analysis of the RNA-Seq assembly problem leads to key algorithmic insights that allowed us to develop Shannon. At a high level, the algorithm has three steps, shown in Figure 3(a): transcript graph construction, copy count estimation and finally estimating transcripts from the graph.

**Figure 3:**
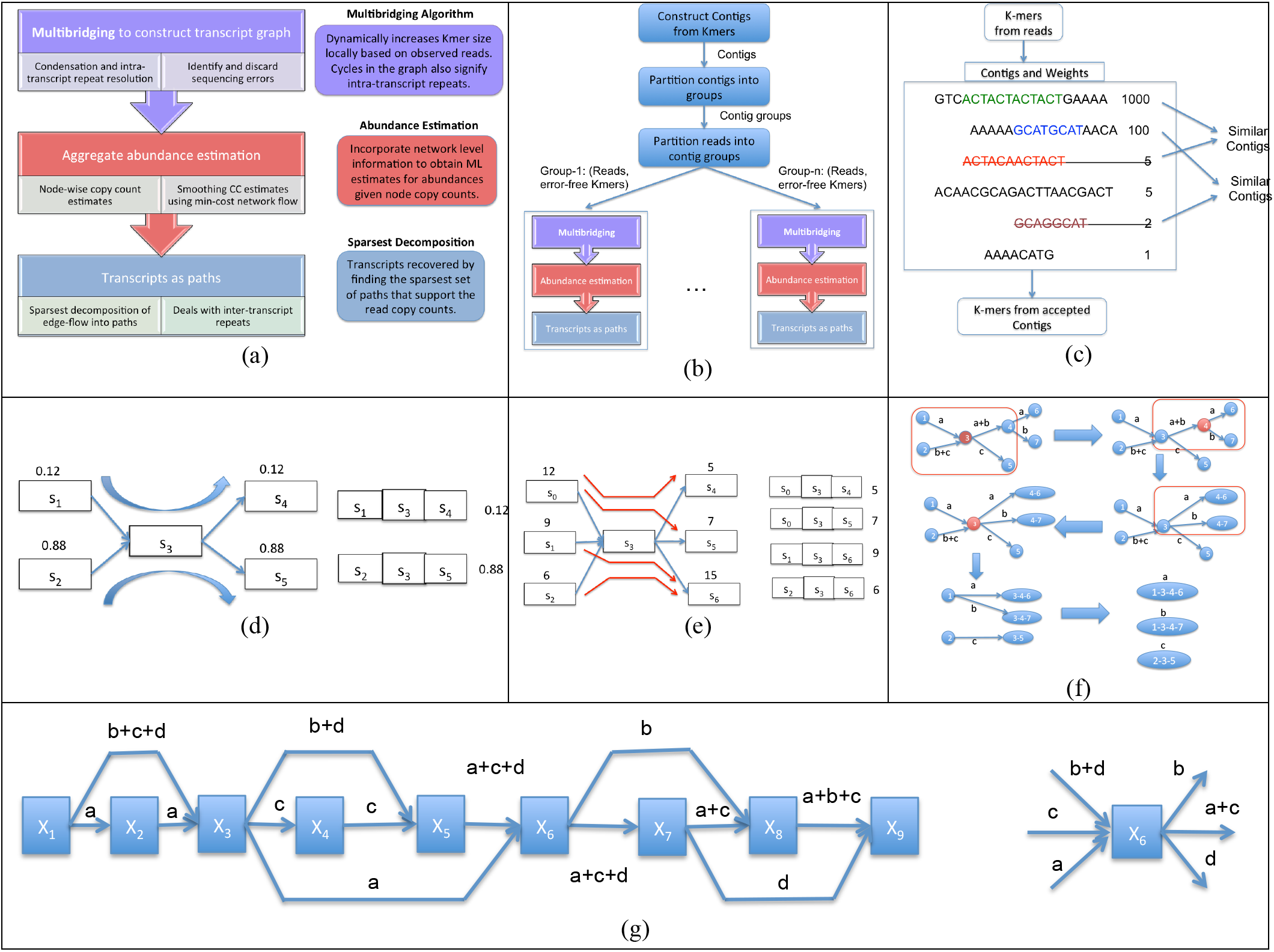
(a) The various steps of the proposed algorithm, (b) Architecture for scaling the proposed algorithm computationally, (c) Structure of error correction algorithm (d) A two-transcript example, where without abundance it is unclear which two paths to reconstruct, (e) A four-transcript example, where the utility of abundance needs well-designed algorithm to find the sparsest flow. In particular, a greedy algorithm will make a mistake and reconstruct a 5-transcript output, (f) A sample run of the algorithm for selecting transcripts from the transcript graph based on sparsest flow using local moves, (g) Example of a real splicing graph from RefSeq PPIP5K1 (Kinase) Gene having 4 isoforms: NM 001190214.1 at abundance a, NM 001130859.2 at abundance b, NM 001130858.2 at abundance c and NM 014659.5 at abundance d. Here nodes *X_1_* correspond to exons, whereas edges and edge-weights correspond to junctions and expected copy counts.

The standard method used by de novo RNA-Seq assemblers is to construct a de Bruijn graph using k-mers of a certain size, and we follow the same first step. However, intra-transcript repeats of length larger than size k but smaller than read length L, which have the potential to be resolved, are left unresolved by the de Bruijn graph. In order to resolve all these repeats in an information-optimal way, we utilize an algorithm called multi-bridging, which is shown to resolve nearly all information-theoretically resolvable intra-sequence repeats.^11^ This algorithm iteratively uses reads to resolve larger and larger repeats, by dynamically and locally increasing the k-value based on the coverage of each node (see Algorithm 3 in Supplementary Methods). Therefore, higher values of k are used in higher coverage regions to assist in resolving repeats, whereas in lower coverage regions small values of k are used so that graph connectivity is maintained. This method is particularly helpful for RNA-Seq assembly where varying abundances among transcripts implies that no single k value may be optimal for the entire transcript graph. In the graph, the copy counts of the various nodes are also stored, using either the average k-mer counts or using the number of reads covering the node (two options in the assembler).

Optionally, the copy-count values can be smoothed using a network flow algorithm to ensure that they constitute a flow (see Supplementary Methods, Section 5), i.e., the sum of copy-counts of all edges into a node is equal to the sum of copy counts of edge going out of the node.^13^, ^14^ In our implementation we did not find significant gains due to this smoothing and therefore it is turned off by default.

Multi-bridging resolves intra-transcript repeats but inter-transcript repeats, comprising of shared exons or duplicated gene segments, remain largely unresolved. From our information-theoretic analysis, it is evident that an information-optimal algorithm should use abundance information carefully. Therefore, we ask for the fewest number of transcripts that best explain the transcript graph and the copy-counts on the graph; this problem can be related to the sparsest flow decomposition problem in graph theory. Indeed, if we do not utilize the copy-count information in our formulation, it may lead to incorrect answers. One such example is shown in Figure 3(d), where there are two possible transcriptomes, which both give rise to the same graph (note that this figure is analogous to Figure 2(b)). By exploiting abundance information, Shannon can uniquely determine the transcriptome. For this simple example, one can exploit the fact that the abundance in the incoming and outgoing edge are the same; we term this the continuity-of-abundance heuristic. General instances of this problem, however, are not solvable by this heuristic; for example, consider Figure 3(e) where there are three incoming and outgoing edges created by four transcripts. The simple greedy approach of matching up the edge-weights does not work in this setting. However, our algorithm still solves this problem correctly.

In general, the sparsest-flow problem has been shown to be NP-Hard (i.e., there is likely no polynomial time algorithm which solves every single instance of this problem), and this has been observed in other reference-based RNA-Seq papers,^13^, ^14^ thus necessitating heuristics such as greedy removal of heaviest paths. This heuristic has also been used in the recent reference-aided assembler Stringtie.^15^ Other ideas for path-selection in reference-aided assembly include minimal path-set incorporating a continuity-of-abundnace heuristic in Cufflinks^16^ and a *l*_1_ minimization algorithm proposed in Isolasso (which also has computational disadvantages, since there can be exponentially many paths, in addition to the fact that it has no informational optimality). Similar heuristics are also employed in de novo assemblers TransAbyss^4^ and Oases.^2^

We propose a novel iterative algorithm that can find a sparse solution to the flow decomposition problem in linear-time (see Supplementary Methods, Algorithm 5 for a description). This iterative algorithm is guaranteed to give the sparsest solution if the abundances are generic (i.e., they are diverse) and a certain technical condition on the graph is met, which is similar to requiring that there are no information bottlenecks (see Figure 2c for an example bottleneck, and Supplementary Materials, Section 4 for more details). Thus, the instances that do not have an information bottleneck are also computationally-efficiently solvable - revealing how information theory overcomes the NP-hardness of the computational problem by focusing only on the relevant instances of the problem. This iterative algorithm, forms the backbone of the transcript-path-selection algorithm in Shannon (see Figure 3(f)). In fact, even for toy examples of multi-isoform genes, existing algorithms fail to tease apart the correct isoforms, whereas Shannon succeeds (see Figure 3(e)). A real example of a kinase gene is shown in Figure 3(g), where Shannon succeeds but existing algorithms fail.

While we described the architecture till now in the absence of errors, we can employ a read error-correction software package before running our assembler to deal with errors. Standard algorithms for DNA read error-correction, for example Quake,^17^ discard kmers that do not have sufficient copy-count and therefore do not work well for RNA-Seq data, which has widely varying abundances. There have been some approaches to read error correction for varying coverage data.^18^, ^19^ We found Quorum^20^ worked best as the pre-error correction stage in our pipeline. This architecture, with a pre-error correction step, works well for small problem instances, however when dealing with large datasets, some modifications are needed to make it efficient in practice (see Figure 3(b)). In order to make Shannon robust in the presence of read errors, we developed a novel error correction algorithm based on another principle from information theory, called successive cancellation (see Online Methods). We first reconstruct representatives of higher abundance isoforms and then remove erroneous contigs that look similar to these higher abundance isoforms, thus eliminating errors that arise from highly abundant isoforms (see Figure 3(c).) Shannon then clusters these contigs into similar groups based on an approximate graph-partitioning algorithm, so that each cluster can be solved independently. The reads and corrected kmers are then assigned to these clusters, so that each cluster now has a smaller set of reads and corrected kmers and the assembly problem corresponding to each cluster can be solved by a separate instance of Shannon in parallel. This error correction and clustering step enables Shannon to run efficiently on large RNA-Seq datasets.

De novo RNA-Seq transcriptome assembly is an important problem with application to both basic biology and biomedicine. As single-cell RNA-Seq assays are improved, higher throughput will allow for assembly of individual cellular transcriptomes,^21^ a task for which Shannon will be well suited. Similarly, while just in its infancy, de novo assembly of cancer transcriptomes will allow for the accurate identification of tumors^22^ and Shannon’s ability to resolve transcripts with overlapping sequence should help to distinguish genes from their fusions. In addition, Shannon should be useful for transcriptome assemblies of non-model organisms whose genomes and transcriptomes are frequently highly repetitive due to whole genome duplication as well as transposon activity.

In addition to its practical value for RNA-Seq assembly due to improved accuracy over previous methods, the information theoretic approach should be useful in solving related problems, e.g. metagenome and meta-transcriptome assembly. Combinatorial optimization formulations of many such problems are NP-hard, but just like in the RNA-Seq assembly problem, information theory can be used to identify problem instances that are both meaningful to solve and efficiently solvable. Detailed exploration of such applications is beyond the scope of this letter, but all our software is freely available and methods are fully described in the Supplementary Materials.

## Author’s contributions

SK, LP and DNT conceived the project and developed the theory and algorithms. SK led the development of the software tool. JH and KM assisted in the software development and algorithmic optimization. SK, LP and DNT wrote the manuscript.

## Acknowledgements

We thank Nicholas Bray for many fruitful discussions. SK was supported in part by the Center for Science of Information, an NSF Science and Technology Center, under grant agreement CCF-0939370, a gift from Qualcomm Inc, and by NIH under award number R01HG008164. JH and KM are supported by the Center for the Science of Information. DNT is supported in part by the Center for the Science of Information and in part by the National Human Genome Research Institute of the National Institutes of Health under award number R01HG008164.

## 1 Methods

### 1.1 Performance evaluation

To evaluate the performance of various assemblers, we consider datasets that have both short reads and long Pacbio reads and follow the methodology described as follows (see Supplementary Figure 1). The short reads are assembled by all the competing assemblers (note: the assemblers do not have access to the long PacBio reads). In both the datasets of interest, the PacBio reads have been used by prior work, potentially in conjunction with the short reads to enhance / augment a standard transcript annotation like RefSeq or Gencode. The number of transcripts reconstructed at more than 90% from this list is shown in Figure 1(a) and Figure 1(e).

The short reads are then used to find transcripts in the enhanced annotation, which are fully covered by kmers in the short reads to get the oracle set. This oracle set is then quantified using abundance estimation software, (we used eXpress^23^) to get the estimated coverage levels of various transcripts. A transcript is said to be reconstructed by an assembler if it has greater than 90% similarity to a reconstructed transcript as measured by BLAT score.^24^ The sensitivity of the reconstruction is then plotted as a function of the coverage levels in Figure 1(b) and Figure 1(f).

The complexity of repeat structures either due to alternative splicing or due to paralogous genes influences the ability of an assembler to reconstruct transcripts. In the simplest case, if a particular transcript shares no kmer with any other transcript in the ground truth, then that transcript should be easy to reconstruct. In order to capture this effect, we define an equivalence class of transcripts, where two transcripts are in the same equivalence class if they share a kmer and thus potentially confusable by short read assemblers. We call two transcripts in the same equivalence class as quasi-forms (similar to isoforms), and segregate the reconstruction performance by the number of quasi-forms in the equivalence class of the transcript in Figure 1(c) and Figure 1(g).

Since the oracle set (or the enhanced annotation) transcripts may not represent an exhaustive set of transcripts in the sample, we cannot assess false positive rates relative to this set. Therefore we utilize the long reads to calculate the false positive rate. A transcript is said to be not false positive, if it is contained inside or contains at least one long read (with 90% similarity score as measured by BLAT). The latter condition is necessary since some of the longer transcripts may not be covered fully by a long read. The fraction of output transcripts which are false-positives is plotted for each software in Figure 1(d) and Figure 1(h).

The plots for the second dataset have two changes: Oases is not included since it crashed every time we ran the software; the memory was seen to climb steadily (utilizing over 90% of the 384GB RAM in the system). The second change is in the methodology to find false positives: We first found that the false positive rate was high for all assemblers - even in this case, Shannon performs the best among all assemblers. The reason for the high false positive rate is the process used to generate Pacbio reads biases against short transcripts (of length lesser than 1000). To quote,^7^ “however, genes with a CSMM only very rarely represented genes smaller than 1 kb, which is likely due to the use of magnetic beads in the loading procedure, which disfavor short fragments.” Therefore to get a fair representation of performance, we only measure false positive rates among reconstructed transcripts of length greater than 1000 in this dataset. To see evaluation on a simulated dataset, see Supplementary Figure 2.

### 1.2 Information Theory

In this paper, we extend the theory of DNA assembly^11^ to the problem of RNA assembly. Compared to DNA assembly, there are two aspects specific to RNA assembly. First, while each transcript can have repeats like a genome does, repeats across transcripts, in the form of shared exons between alternately spliced isoforms, are typically much longer and causes the information bottleneck for RNA assembly. Second, unlike the repeats in the DNA assembly problem, these inter-transcript repeats can be resolved even when the reads are shorter than the shared exon length. This is achieved by exploiting the fact that different transcripts typically have widely varying abundances. An example of such a resolvable repeat pattern is shown in Figure 2(b). Thus, varying abundances, which seem to complicate the RNA assembly problem, can actually be beneficial. To establish the limits to RNA assembly, our theory characterizes all the bottleneck intra-transcript repeat patterns that cannot be resolved even with the help of varying transcript abundances, and gives an algorithm that can resolve most other repeat patterns (Supplementary Materials). An example of an unresolvable repeat pattern is shown in Figure 2 (c).

By characterizing the bottleneck repeat patterns, our theory computes tight upper and lower bounds on the critical read length /_crit_ needed to reconstruct a transcriptome. The theorem for information-theoretic reconstruction is given below.

#### Theorem 1.

(**Information theoretic condition**) The following condition is necessary for unique reconstruction of a transcriptome T from reads of length L by any algorithm:

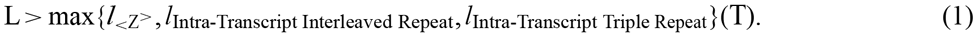

The proposed iterative algorithm reconstructs the transcriptome uniquely, assuming generic abundances, if

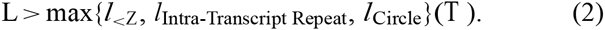

Here the various quantities on the RHS correspond to the maximum size of repeats with certain structures, either inside transcripts or across transcripts (we refer the interested reader to the Supplementary Material, Section 3, for definitions of these quantities). Among all the bounds on the read-length, the biting-condition in biological samples we have tested on, comes from the *l*_< z_ in the necessary condition and *l*_< z_> in the sufficient condition; and these two numbers evaluate to the same quantity, implying that the two bound match, yielding the critical read length. For example in a reference mouse-transcriptome, *l_crit_* = 4077. This long critical read length is due to the existence of a few complex isoforms in the transcriptome. Giving up on recovering these few complex isoforms, the rest of the transcriptome can be reconstructed with potentially smaller read length. More generally, our characterization of the bottleneck repeat patterns leads to information theoretic bounds on the fraction of a given transcriptome that can be reconstructed (under the idealistic setting of infinite coverage depth and error-free reads). Figure 2(d) shows these bounds for the HESC transcriptome. Assembly algorithms that do not use abundance information cannot resolve repeat patterns like the ones shown in Figure 2(b) and thus cannot recover as many transcripts as the information theoretic bounds suggest. Our theory can calculate a bound on the maximum fraction of transcripts these algorithms can reconstruct; this bound is shown as the lower curve in Figure 2(d) for the HESC transcriptome. The large gap between this bound and the information theoretic bound is the theoretical basis for the success of Shannon on real data as shown earlier.

### 1.3 Assembler Design

The key algorithmic insights underlying the assembler arise from the information theoretic analysis of the RNA-Seq assembly problem. At a high level, the algorithm has three steps, shown in Figure 3a: transcript graph construction, copy count estimation and finally estimating transcripts from the graph. In this description, we will first describe the algorithm in the error-free case and then describe the methods to deal with errors and how to scale the algorithm.

We have key innovations in all three steps, which we sketch below.

- Constructing the transcript graph: A typical first step required in de novo transcriptome assembly from short reads is the construction of a k-mer (DeBruijn) graph: a fixed length k is chosen and reads are chopped into subsequences of size k, called k-mers. To count k-mers we use a fast parallel k-mer counting software package called Jellyfish.^25^ A graph is constructed in which the nodes are k - 1-mers and two nodes are connected if the k - 1 mer corresponding to the nodes overlap head-to-tail by k - 2 and the (k + 1)-mer formed by appropriate concatenation is present in the reads. This graph is condensed by collapsing simple chains of nodes. The problem of choosing an appropriate value of k is not an easy one, since, in the context of RNA-seq assembly (as opposed to say DNA assembly), the abundances take values in a large range. If k is chosen too small, then transcripts which share larger common sequences can be confused; whereas if k is too large, then the graph will become disconnected for low-abundance transcripts. Thus no particular value of k may be optimal uniformly throughout the graph and different values of k may need to be chosen for different nodes in the graph. A key innovation in this direction came in an earlier work on information-theoretic DNA assembly,^11^ where an algorithm called multi-bridging was proposed. In Multi-bridging, the K-mer size is chosen dynamically based on the quantity and sequence of reads supporting the different portions of the graph. For computational efficiency, the algorithm starts with a certain k-value and increases it dynamically in the different portions of the graph. The algorithm also brings back the reads supporting each node in a careful manner in order to maximize reconstruction probability. While maintaining the DeBruijn graph, the copy-counts of the nodes and edges are also maintained. For details of the multi-bridging algorithm, we refer the reader to Algorithm 3 in Supplementary Material. In the presence of errors, standard techniques for DeBruijn graph trimming in the presence of errors are used. In particular, iteratively, the algorithm searches for pairs of nodes that are connected by two possible paths of length 1, i.e., having exactly one intermediate node in each path. If the sequences generated by the two paths are similar, then the path with lower abundance is rejected. Another iterative error-correction algorithm is applied, which rejects nodes that have copy count lesser than a threshold or size lesser than a threshold or if a node is a hanging node (has no outgoing or incoming edge) or if a node is dispensable (a node is said to be dispensable if each of its out-nodes has in-degree strictly larger than 1 and each of its in-nodes has out-degree strictly larger than 1. Notice that nodes created by typical errors will be dispensable). Once a node is rejected, the algorithm locally recondenses the graph - so that potentially larger nodes can be created by collapsing new simple chains induced by error correction. Once the reads are brought back into the graph via multi-bridging, they are used to resolve certain patterns of repeats. However, some of them may not be fully exploited by multi-bridging; in our algorithm, we write out such nodes into a file of known-paths, i.e., these are paths in the graph that are supported by reads. In the presence of paired-end reads, after multi-bridging, the algorithm searches for paths of length shorter than the insert-size within which paired-end reads are placed on the graph. If the path between the paired-end read is unique, then the algorithm treats this as a longer read of size equal to the fragment size. Thus, in Shannon-RNA, paired-end reads provide an advantage over single-end reads in terms of repeat resolution.
- The copy count of the k-mers associated with each node yields an estimate of the aggregate abundance of all the transcripts containing that exon. An alternative mode in our assembler uses the reads placed back on the graph in order to estimate the node / edge copy-counts more accurately. However, in either mode, these estimates are noisy due to the finite coverage and varying abundance of the transcripts. An innovation in the current algorithm is the utilization of a smoothing step that takes the estimated coverage and calculates the smoothed coverage using either a maximum likelihood or an *l_1_* regularizer. This mode is currently optional and can be enabled as required. (We refer the reader to Supplementary Material Section 5 for details.)
- In the final step, the algorithm needs to return a subset of paths through the graph in order to reconstruct the transcripts. This corresponds to the mathematical problem of finding the sparsest flow through the network that explain the smoothed copy counts. The sequences of these paths are output as transcripts and their flow values correspond to the transcript abundances. However, prior work^14^ has noted that the network sparsest flow problem is NP-hard, and therefore resort to heuristics. The main idea in our work is to derive a novel linear time algorithm, which can solve a family of instances of the problem efficiently, which includes most biologically relevant instances. The algorithm proceeds by first 1) decomposing the global problem into many local sparsest flow problems, one for each node and then 2) solving each local sparsest flow problem efficiently using ideas from sparse-signal recovery theory, using a randomized *l*_1_-relaxation. This algorithm is not only computationally efficient, but also is able to reconstruct provably correctly beyond the minimal information threshold. We refer the reader to the Supplementary Material (Algorithm-5 and Section 4). An example of a two-transcript system, where abundance is needed for successful reconstruction is shown in Figure 3d. Another example of a four-transcript system, where abundance is needed for successful reconstruction and furthermore, greedy algorithms for abundance based reconstruction^13^, ^14,15^ cannot reconstruct accurately. Finally, an example run of the algorithm is shown in Figure 3f, where the algorithm starts to decompose node 3, is unable to find a unique sparsest path locally and therefore moves on to node 4 which it successfully resolves before returning to resolve node 3 successfully.

#### 1.3.1 Dealing with errors

To deal with errors, we have a pre-error correction stage where reads are fed to error correction package Quorum^20^ - this package uses quality scores in addition to kmer contiguity to correct errors. We also have an additional in-house error-correction step to ensure that the deBruijn graph algorithm that follows, runs smoothly. Popular methods for k-mer error correction in DNA sequence, such as Quake,^17^ utilize thresholds on k-mer copy counts (or their appropriately quality-weighted analogues) in order to determine which k-mers are true and which are erroneous. However, the large-range of abundance values present in RNA-Seq data preclude the utility of such methods - since no appropriate threshold can be found. Therefore, we propose a new contig-based error correction algorithm. While this algorithm is inspired by the algorithm for constructing contigs in Trinity, there is a critical distinction - our algorithm is based on a method in information theory called successive cancellation.

We propose the following algorithm, shown pictorially in Figure 3c. We first sort the list of k-mers according to abundance and then start with the heaviest k-mer, find the heaviest k-mer that overlaps for k - 1 bases with the present k-mer and use that to extend the contig, till no further extension is possible. This process is repeated to build a list of contigs, while removing the k-mers once used from further consideration. For each contig, we also maintain the average k-mer weight. The contigs thus constructed have good representative contigs as well as erroneous contigs. We remove contigs, which are very similar to another contig of higher weight. This process is called successive cancelation, and traces back to ideas in multi-user communication in information theory.^26^ After successive cancellation, we further reduce the number of contigs by keeping only contigs whose average weight and length are large enough to satisfy a certain relationship between them. At the end, from the contigs that are accepted, we list down the k-mers and these k-mers are returned as the accepted k-mers, to be used for constructing the initial De-Bruijn graph for the multi-bridging algorithm.

#### 1.3.2 Scaling the Algorithm

Having done error correction, it is possible to run the proposed algorithm for the error-free case. However, in order to parallelize the remainder of the steps, we propose to perform an additional step to decompose the problem into smaller sub-problems that can then be solved independently and parallely. In order to accomplish this, we have to partition the list of k-mers into groups, and furthermore, we have to assign reads into these groups as well. Our method for decomposing the problem comprises of decomposing the set of contigs into groups. First we construct a graph, where contigs are nodes and edges between contigs mention the number of k-mers shared between the two contigs. We would like to partition the graph into clusters such that the weight of the edges cut by the k-mers is minimized. This problem is called approximate graph partitioning, and while the problem is NP-hard in the worst case, there are excellent implementations of algorithms available that work very well in practice. We propose to use METIS,^27^ which is a software package for approximate graph partitioning. The output of METIS is a clustering of contigs. The k-mers in the contigs are naturally partitioned into various clusters. In order to partition the reads, we look at how many k-mers are shared with any particular cluster and assign the read to the cluster to which it shares the maximum number of k-mers. At the end of this process, we have a set of smaller problems, each comprising of a list of cleaned-up k-mers and a list of reads. Now each of these problems can be solved independently and parallely using the algorithm in Figure 3a. We use GNU parallel^28^ in order to solve this trivially parallelizable problem. While most of the transcripts are resolved by this method, transcripts that have straddled between distinct clusters and that got cut off due to the approximate graph partitioning will be missed in the output. Therefore, we do another partitioning step, where the parameters are adjusted so that edges cut in the first partitioning are weighted higher, so that they are less likely to be cut again. The algorithm is then run parallely on all the instances, including the repartitioned instances. This gives rise to an increased sensitivity of the algorithm. It is to be noted that this process does not give rise to false positives - since paths that arise in the repartitioning are likely to be subpaths of the true paths (which may have been decoded in the first run) - and if needed, they can be discarded using a post-processing step.

**Code availability:** Shannon is available as a open-source software that can be downloaded from http://sreeramkannan.github.io/Shannon/

## Footnotes

1 ^a^Shannon is available as an open-source software and can be downloaded from http://sreeramkannan.github.io/Shannon/.

